# Development and evaluation of a formulation of probiont *Phaeobacter inhibens* S4 for the management of vibriosis in bivalve hatcheries

**DOI:** 10.1101/2022.12.27.522043

**Authors:** Evelyn Takyi, Jason LaPorte, Saebom Sohn, Rebecca J. Stevick, Erin M. Witkop, Lauren Gregg, Amanda Chesler-Poole, Jessica Small, Meredith M. White, Cem Giray, David C. Rowley, David R. Nelson, Marta Gomez-Chiarri

## Abstract

Larval eastern oysters (*Crassostrea virginica*) grown in shellfish hatcheries are susceptible to bacterial diseases, particularly vibriosis. Probiotics are microbes that confer health benefits to the host and have been identified as promising tools to manage diseases in aquaculture. The marine bacterium *Phaeobacter inhibens* S4 (S4) protects larval eastern oysters against challenge with the bacterial pathogen *Vibrio coralliilyticus* RE22 (RE22). A concentrated liquid formulation of probiont S4 that maintained high cell viability after long-term storage was developed for commercial use in shellfish hatcheries. The safety and efficacy of the formulation was tested in six different trials in two hatcheries. The S4 formulation was added to *C. virginica* larvae culture tanks daily at 10^4^ colony forming units (CFU)/mL from day 1 post fertilization until day 6, 12, or 14, depending on the trial. Treatment of larvae in the hatchery with the S4 formulation did not significantly affect the survival and growth of the larvae. Formulated probiont S4 treatment in the hatchery led to a significant increase in Relative Percent Survival (RPS) when larvae were subsequently challenged with the pathogen RE22 (10^5^ CFU/mL) for 24 hours in a laboratory challenge, as compared to probiotic-untreated RE22-challenged larvae (Relative Percent Survival increase of 46 - 74%, *p* < 0.05). These results suggest that this novel S4 formulation is a safe, easy to use, and effective tool in preventing larval losses due to vibriosis in hatcheries.

## 1. Introduction

The eastern oyster, *Crassostrea virginica*, is a bivalve species with significant ecological and economic importance to the Gulf of Mexico and Atlantic coastal communities of North America [1, 2]. Oyster production through aquaculture in the United States totaled 219 million dollars (USD) in 2018 [3]. Hatchery production of oyster seed is crucial for ensuring a constant and sufficient supply of juveniles to support the oyster industry. However, changes in environmental conditions and disease outbreaks are limiting factors for the growth of aquaculture production [4]. Vibriosis, a disease caused by pathogenic bacteria in the genus *Vibrio*, has been an issue of particular concern in bivalve hatcheries. Various strains of *Vibrio* spp. that are pathogenic to oyster larvae lead to a rapid and high rate of larval mortality in hatcheries, resulting in substantial economic loss to the oyster industry [5,6,7,8]. Techniques for managing disease outbreaks in hatcheries include the use of water treatment systems (*e.g*., filtration, ultraviolet light, pasteurization) and labor-intensive biosecurity measures (*e.g*., cleaning of equipment) to avoid the introduction and spread of pathogens [5]. Antibiotic usage in bivalve shellfish hatcheries is discouraged because of the potential development of resistance by bacteria and negative impacts on healthy oyster microbiota [9,10]. Despite significant efforts to treat the water supply, pathogenic vibrios are still detected in shellfish hatcheries and may cause shellfish mortality in opportunistic conditions [5].

The use of probiotics has emerged as a potential tool to reduce mortalities in the rearing of aquatic organisms and manage disease outbreaks in aquaculture [11,12,13,14]. Probiotics are defined as live, non-pathogenic microorganisms which, when administered in adequate amounts, confer a health benefit to the host [15]. In aquaculture, probiotics are administered as either food supplement or as an additive to the water [16,17,18,19]. Candidate probiotics for use in invertebrate aquaculture include a variety of gram-negative and gram-positive bacteria, yeast, and unicellular algae. Depending on the probiotic species used, these health benefits are derived from a variety of complementary mechanisms including improvement of water quality, enhancement of host nutrition through the production of supplemental digestive enzymes, competition with pathogenic bacteria, production of antimicrobial compounds, host immunomodulation, and modulation of microbial community structure to promote health [11,20,21,22,23,24].

The marine bacterium *Phaeobacter inhibens* S4 (S4) is a gram-negative alpha-Proteobacterium in the *Rhodobacter* clade. Several *Phaeobacter* species exhibit inhibitory activity against a wide variety of marine pathogens such as *Vibrio coralliilyticus* RE22, *Vibrio anguillarum, Vibrio tubiashii*, and *Aliiroseovarius crassostreae* [25,26,27,28,29] and have been shown to effectively colonize surfaces by forming dense biofilms [30]. Previous studies have also demonstrated the probiotic ability of probiont S4 to prevent larval eastern oyster mortality against bacterial infection in laboratory and hatchery experiments [28,31]. Mechanisms of S4 protection include biofilm formation, secretion of the antibiotic tropodithietic acid (TDA), quorum quenching by which S4 represses gene expression of virulence factors in the shellfish pathogen *V. coralliilyticus* RE22, and host immune modulation [22,30,32].

Although probiont S4 demonstrated promising results for limiting *V. coralliilyticus* infections in bivalve aquaculture hatcheries, the probiont needs to be delivered daily to be effective [31,33], and daily preparation of fresh cultures in the hatchery is impractical (personal communication with hatchery personnel). A standardized, stable, commercially-produced formulation of probiont S4 would offer advantages such as ease and convenience in storage, handling, and delivery at the hatchery. Commercially formulated probiotics mostly include dry products such as wettable powders, dusts, granules, and liquid products such as cell suspensions in water, oils, and emulsions [11]. Most commercial probiotics available in the market for aquaculture are formulated from a mixture of gram-positive bacteria that show high survival after freeze drying. Examples include Prosol (*Bifidobacterium longum, Lactobacillus acidophilus, Lactobacillus rhamnosus, Lactobacillus salivarius, Lactobacillus plantarum*), Engest Probiotics (*Bacillus subtilis, Bacillus licheniformis, Bacillus megaterium*) for shrimp [34,35] and Bioplus (*B. subtilis, B. licheniformis*)for rainbow trout [36]. To the best of our knowledge, the only gram-negative bacteria commercially formulated is Eco-Pro (*Rhodopseudomonas palustris*), used for improving water quality in ponds [37].

Based on previous research showing the efficacy and safety of using daily treatments of freshly grown cultures of *Phaeobacter inhibens* S4 as a probiont in bivalve larval culture, both in laboratory and hatchery experiments [28,31,33], the present study developed a novel liquid formulation for the gram-negative bacterium S4, allowing for ease of routine application in the hatchery at a commercial scale. This novel liquid formulation was tested for its safety, efficacy, host protection, ease in handling, and delivery in bivalve hatchery facilities. The results demonstrate that the formulation showed similar performance in the hatchery as previously reported for freshly cultured S4 [31,33] and pre-treatment in the hatchery consistently protected eastern oyster larvae from experimental challenge with the bacterial pathogen *V. coralliilyticus* RE22.

## 2. Materials and Methods

### 2.1. Bacterial strains

Bacterial strains *Phaeobacter inhibens* S4Sm (probiont, named S4 thereafter) and *Vibrio coralliilyticus* RE22Sm (pathogen, RE22) (both are streptomycin resistant strains by spontaneous mutation) were maintained as stocks in 50% glycerol at −80°C until use. Bacteria were cultured on yeast peptone with 3% sea salt (mYP30) media (5 g /L of peptone, 1 g/L of yeast extract, 30 g/L of ocean salt (Red Sea Salt, Ohio, USA)) at 27°C with shaking at 175 rpm as described in [28] unless otherwise indicated.

### 2.2. Development of a liquid probiotic formulation for *Phaeobacter inhibens* S4

Initial trials in the development of spray dried or freeze-dry (lyophilized) formulations for *Phaeobacter inhibens* S4Sm using mannitol or sucrose were not successful [38]. Therefore, a liquid formulation was developed. Bacteria from glycerol stocks stored at −80°C were streaked for isolation on a mYP30 agar plate and incubated at 27°C for 24 - 48 hours. A single S4Sm colony was inoculated into Luria Broth with 3% sea salt (mLB30, pH 7) or mYP30 growth medium and incubated at 27°C with shaking for 48 hours, until reaching stationary phase (~10^9^ CFU/mL). Four different formulation methods were tested for viability after storage for 6 weeks: (1) S4 mLB30 stationary broth undiluted cultures stored without shaking at 4°C (LB_4); (2) S4 mLB30 stationary broth cultures diluted 1:1 with 3% filtered sterile artificial seawater (FSSW) and stored at 4°C (LB_SW_4) or (3) 22°C (LB_SW_22), and (4) S4 mYP30 stationary broth cultures diluted 1:1 with FSSW and stored at 4°C (YP_SW_4). The viability of the formulations was determined at 0, 2, 4, and 6 weeks in each of the storage conditions by spot plating serial dilutions in triplicate on mYP30 agar plates and counting colony forming units (CFU/mL) after 24 – 48 hours of growth [30]. Culture and formulation of *Phaeobacter inhibens* S4 were scaled up for commercial production by Kennebec River Biosciences using proprietary methods for large scale bacterial production and the liquid formulation protocol reported here (*i.e*., dilution of stationary high titer S4 mYP30 cultures 1:1 in FSSW).

### 2.3. Laboratory challenge of larvae treated with freshly cultured or formulated S4 probiotic

Laboratory challenge assays were conducted following protocols described by [28]. Briefly, eastern oyster larvae (7 days post fertilization, 100 – 150 μm in size) were obtained from the Aquaculture Genetics and Breeding Technology Center, Virginia Institute of Marine Science (VIMS) hatchery. Oyster larvae (50-100 per well) were placed into 6-well plates with 5 mL of filtered sterile artificial sea water (FSSW, 28 PSU). Larval oysters were fed with commercial algal paste (20,000 cells/mL; Reed Mariculture Inc., San Jose, CA) prior to addition of probiotics to enhance the ingestion of probiotics. Freshly cultured or formulated S4 were added to larvae in wells designated for each probiotic treatment at a concentration of 10^4^ CFU/mL and incubated at room temperature with gentle shaking. After 24 hours, the pathogen RE22 was added to each well at a final concentration of 10^5^ CFU/mL. Control wells included unchallenged larvae (no S4 or RE22) and larvae incubated with S4 but without the pathogen. Each treatment was run in triplicate. Larval survival was determined 24 hours after the pathogen was added using the neutral red technique [39]. Survival was calculated by using the formula: Survival (%) = 100 x (number of live larvae/total number of larvae). The relative percent survival (RPS) of probiotic pretreated and RE22-challenged larvae compared to the RE22-challenged control was calculated using the formula: RPS (%) = [1 - (% Mortality treatment / % Mortality control)] x 100 [28].

### 2.4. Hatchery trial set up

Hatchery experiments were conducted at the Aquaculture Genetics and Breeding Technology Center, Virginia Institute of Marine Science (VIMS) hatchery (Gloucester, Virginia, USA) and Mook Sea Farms hatchery (MOOK; Walpole, Maine, USA). At VIMS, four independent trials (Trials 1-4) were run with 60 L flat-bottom larval rearing tanks used for each trial (Table 1). Tanks (minimum of 3 per treatment) were randomly assigned to the following treatments: no probiotics (control) or S4 formulation (probiotic treatment). Broodstock were spawned at the hatchery using standard techniques for each trial and experiments were initiated by adding 3.6 × 10^5^ to 6 × 10^5^ larvae (6.2-10 larvae/mL) to each static tank on day 1-2 post fertilization. Larvae in each tank were fed with a hatchery-reared microalgal diet consisting of *Pavlova pingus, Chaetoceros negrocile*, and *Tetraselmis chui*. Trials 5 and 6 were conducted at MOOK. In Trial 5, 5.2 × 10^7^ larvae (17.3 larvae/mL) were raised in each of two single 3000 L static tanks (one control, one treated with S4) from day 1 to day 8 post fertilization, and then larvae from each tank were distributed into 3 × 200 L flowthrough tanks from day 9 to day 12. In Trial 6 at MOOK, larvae were raised in 15 L buckets from days 1 to 12. Larvae in each tank at MOOK were fed with a hatchery-reared microalgal diet consisting of *Pavlova lutheri* (CCMP1325), *Tetraselmis* sp. (CCMP892), *Tisochrysis lutea* (CCMP1324), and *Chaetoceros muelleri* (CCMP1316). Probiotic formulation were added daily at a dose of 10^4^ CFU/mL at the time of algal feeding from day 1 (24 hours post fertilization) until the termination of the trial. The trials ended at day 12 or 14 (immediately prior to larval setting, Trials 1, 3, 5, and 6) or earlier if larval performance was low (6 days, Trial 2) or the hatchery needed the tanks for routine production (7 days, Trial 4). Larval tanks were drained down every other day for size grading of larvae, cleaning of tanks and for maintenance of water quality [40].

**Table 1.**
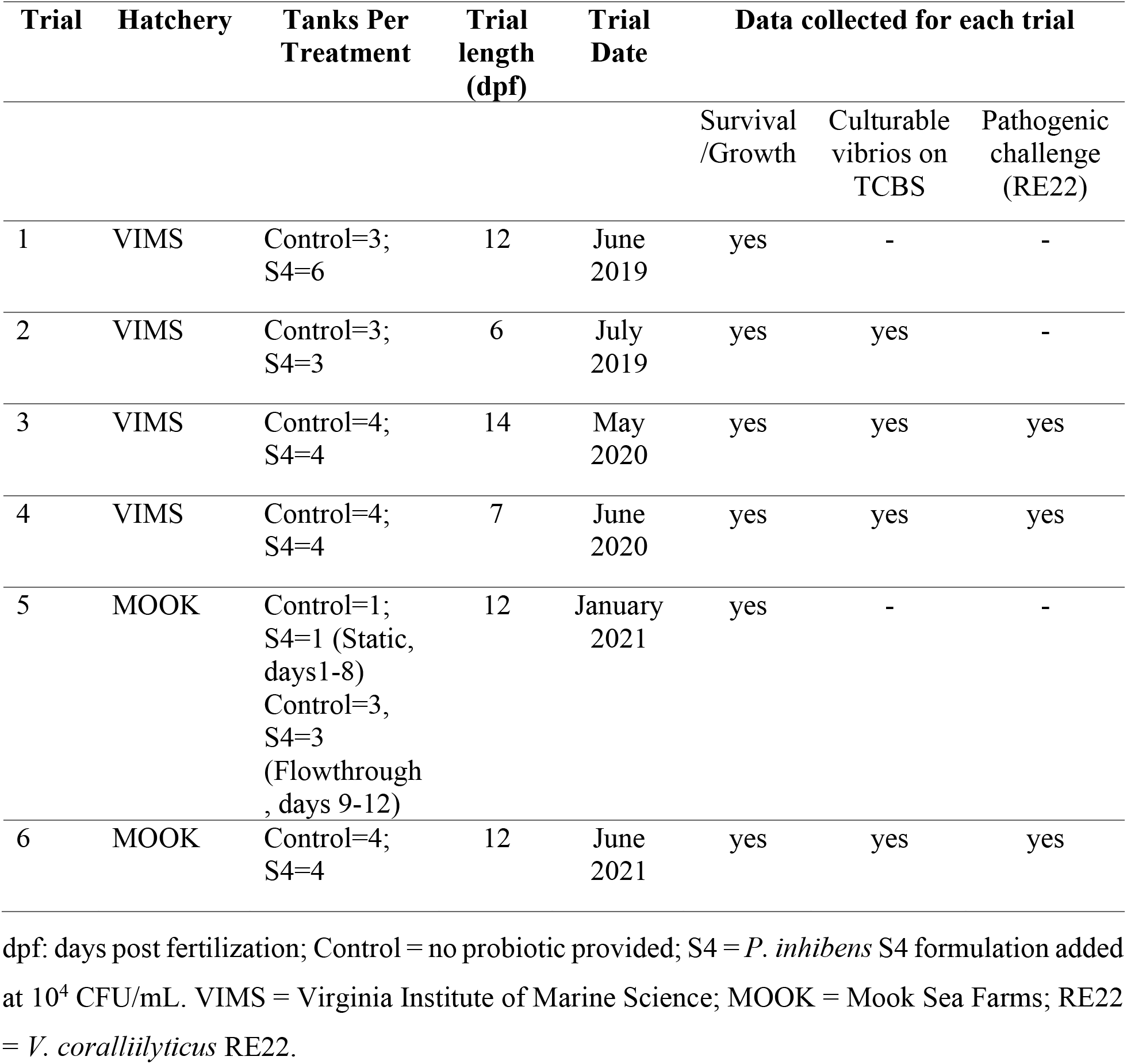
Hatchery trials performed in this study.

### 2.5. Evaluation of the effect of S4 formulation on larval growth and survival during hatchery trials

Data for each trial were collected every 2 days during the trial period at the time of drain down. Larvae from each tank were collected on nylon mesh screens, rinsed, and transferred to 150 L tanks filled with 100 L in trial 5 and 400 mL beakers filled with filtered seawater to 200 mL in trials 1, 2, 3, 4, and 6. At MOOK, after gentle stirring of the water to evenly distribute larvae, a micropipette was used to collect six 1000 μL larval samples into 6 well plates and then immobilized with 70% isopropyl alcohol and counted using a dissecting microscope. Larval sizes were estimated based on the percentage of larvae retained on standard mesh sizes (325, 270, 230, 200, 170, 140, 120, 100, 80, 70, 75; Mook and VIMS) and by image analysis (VIMS only). For image analysis and health assessments, a micropipette was used to collect four 50 μL larval samples. Each sample was placed on a gridded Sedgewick Rafter counting cell installed on the microscope stage. Larvae were initially observed under a 4x objective for motility, overall shape, and gut coloration to make a health assessment and were assigned a health rating from 1 (poor health) to 3 (good health). Larvae were then temporarily immobilized with a 2:1 mixture of freshwater and 70% isopropyl alcohol. Larvae were counted under a microscope and percentage survival was calculated and recorded. Larval sizes were observed under 10x objective magnification and an ocular micrometer was used to measure the longest axis of each larval shell. Larval specific growth rate at the end of each trial was calculated from the larval sizes using the formula: SGR (Specific Growth Rate (% per day)) = ((LnLt - LnLo)/t) x 100 where LnLt = ln final shell length (μm), LnLo = ln initial shell size (μm), and t = time (days) [41].

### 2.6. Determination of levels of *Vibrio* spp. in hatchery larval samples

The total number of culturable *Vibrio* spp. was determined for trials 2, 3, 4, and 6 using a plate count method on thiosulfate-citrate-bile salts-sucrose medium (TCBS, Difco) [31]. Samples were collected from water in the rearing tank (10 mL) and larval oysters (~1,000) during drain down events in the hatchery. Oyster larvae were rinsed with FSSW, homogenized using a sterile pestle, and suspended in 1 mL FSSW. Samples were serially diluted and 10 μL of each dilution were spot plated on TCBS agar plates in triplicate. The inoculated plates were incubated for 16 - 20 hours at 28 °C and colonies were counted. Results were expressed as CFU/mL.

### 2.7. Laboratory pathogen challenge of probiotic-treated larvae from hatchery

Since pathogens could not be introduced into the hatcheries, a subsample of about 1,000 larvae from each tank were collected during drain down events and shipped overnight on ice (kept cool, not frozen) to the laboratory at University of Rhode Island. Oyster larvae (~50-100 per well) were placed in 6-well plates and challenged with the pathogen RE22 at a final concentration of 10^5^ CFU/mL following the methods described in the laboratory challenge section above. Controls included 3 wells of non-challenged larvae per tank and treatment.

### 2.8. Statistical Analysis

All statistical analyses were performed in the R statistical computing environment, version 4.0.2 [42]. Data were checked for normality and homogeneity of variance prior to selection of the statistical method. Each trial was analyzed separately to determine effect of S4 formulation, as compared to non-treated control, on each of the parameters. Larval oyster percent survival was subjected to arcsine square root transformation prior to statistical analysis. Due to large differences in larval performance between some of the trials (see results below), one-way analysis of variance (ANOVA) was used to analyze data for each trial separately and determine significance between treatments within each trial. Tukey’s multiple comparison tests were used for post-hoc pairwise comparisons. A *p*-value ≤ 0.05 was considered to be statistically significant.

## 3. Results

### 3.1. Viability of formulated S4 under various storage conditions

The viability of the different S4 formulation methods described were assessed to determine the concentration and stability of the bacteria over time. The viability was assessed biweekly during storage for 6 weeks (Figure 1). Formulation method (storage media and temperature) had a significant effect on the viability of probiont S4 at the end of the 6 weeks (One-way ANOVA; *p* < 0.05; Figure 1). The formulations in which dense cultures of S4 (grown in either mLB30 or mYP30) were diluted 1:1 in FSSW and then stored at 4°C (LB_SW_4 and YP_SW_4) showed significantly higher viability at the end of the 6 weeks (declines of 0.52 log and 0.6 log) as compared to undiluted cultures stored at 4°C (LB_4) or the diluted cultures stored at 22°C (LB_SW_22) (declines of 1.32 and 1.34 log, respectively). Based on these results, the YP_SW_4 formulation was used in the following hatchery trials.

**Figure 1.**
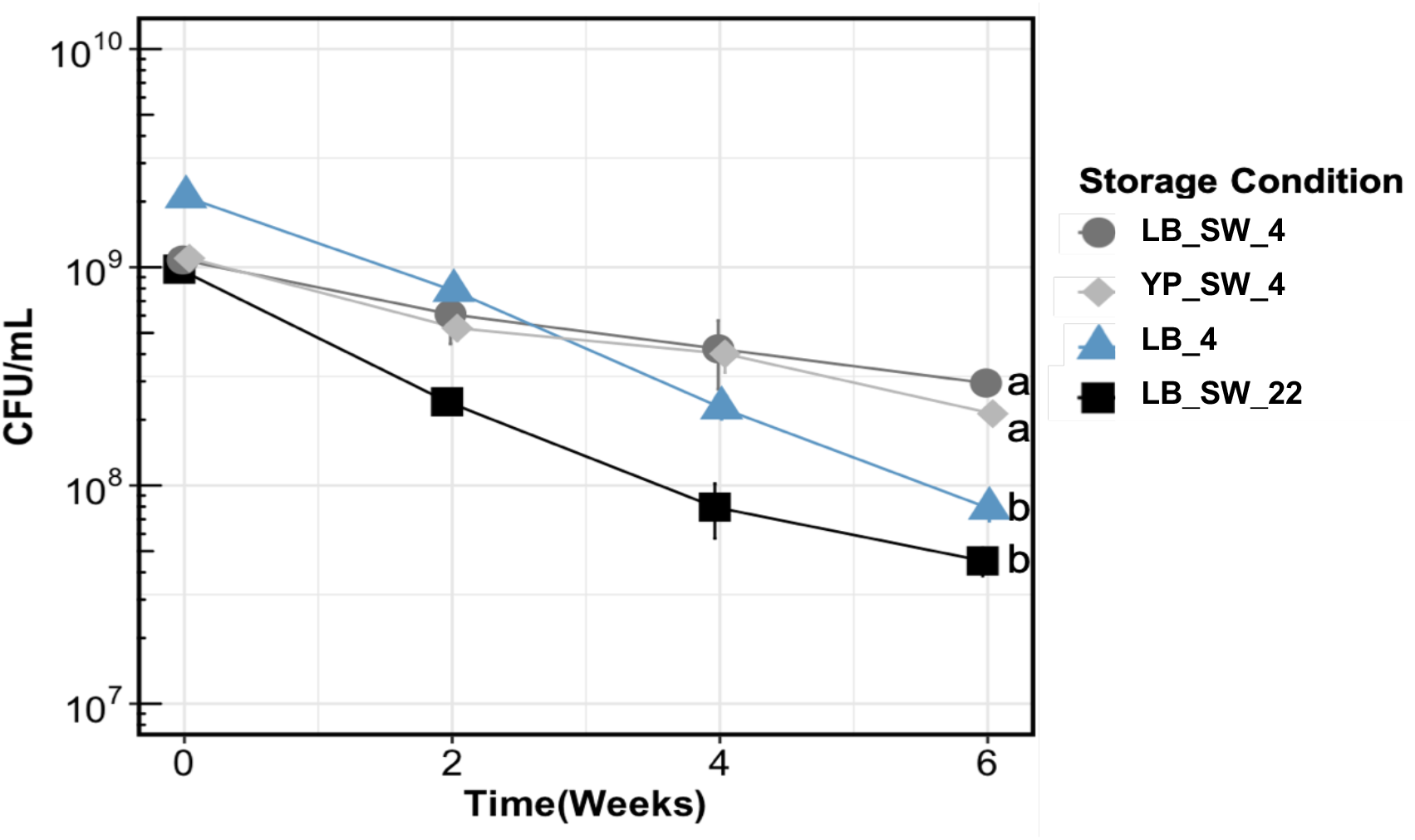
Impact of formulation method on the viability of S4. Bacterial cultures were stored for 6 weeks in four different conditions and sampled biweekly. Data expressed as mean ± SD of CFU/mL of S4 (n=3). S4 = *Phaeobacter inhibens* S4; LB_4: S4 mLB30 undiluted broth culture stored at 4°C; LB_SW_4: S4 stored in diluted mLB30 broth (1:1 with FSSW) at 4°C; LB_SW_22: S4 stored in diluted mLB30 (1:1 with FSSW) and stored at 22°C; YP_SW_4: S4 stored in diluted mYP30 broth (1:1 with FSSW) and stored at 4°C; FSSW: Filtered Sterile Seawater. Different letters indicate statistically significant differences based on Tukey’s pairwise comparisons (Oneway ANOVA, *p*<0.05; for samples on week 6).

### 3.2. Fresh and formulated S4 protected larvae against pathogen challenge in the laboratory

Pretreatment of larvae with freshly cultured S4 or formulated S4 (YP_SW_4) had no detrimental effect on larval survival (*i.e*., in the absence of pathogen challenge) over a 48 hour period (Figure 2). Challenge of untreated control larvae with the pathogen RE22 led to significant larval mortality (90% decrease, One-way ANOVA on transformed survival data, *p* < 0.05). Pretreatment with either the freshly prepared or the formulated S4 significantly increased larval survival following RE22 challenge as compared to non-treated controls (40% - 60%; *p* < 0.05) and the levels of survival did not differ between the two (One-way ANOVA;*p* > 0.05; Figure 2).

**Figure 2.**
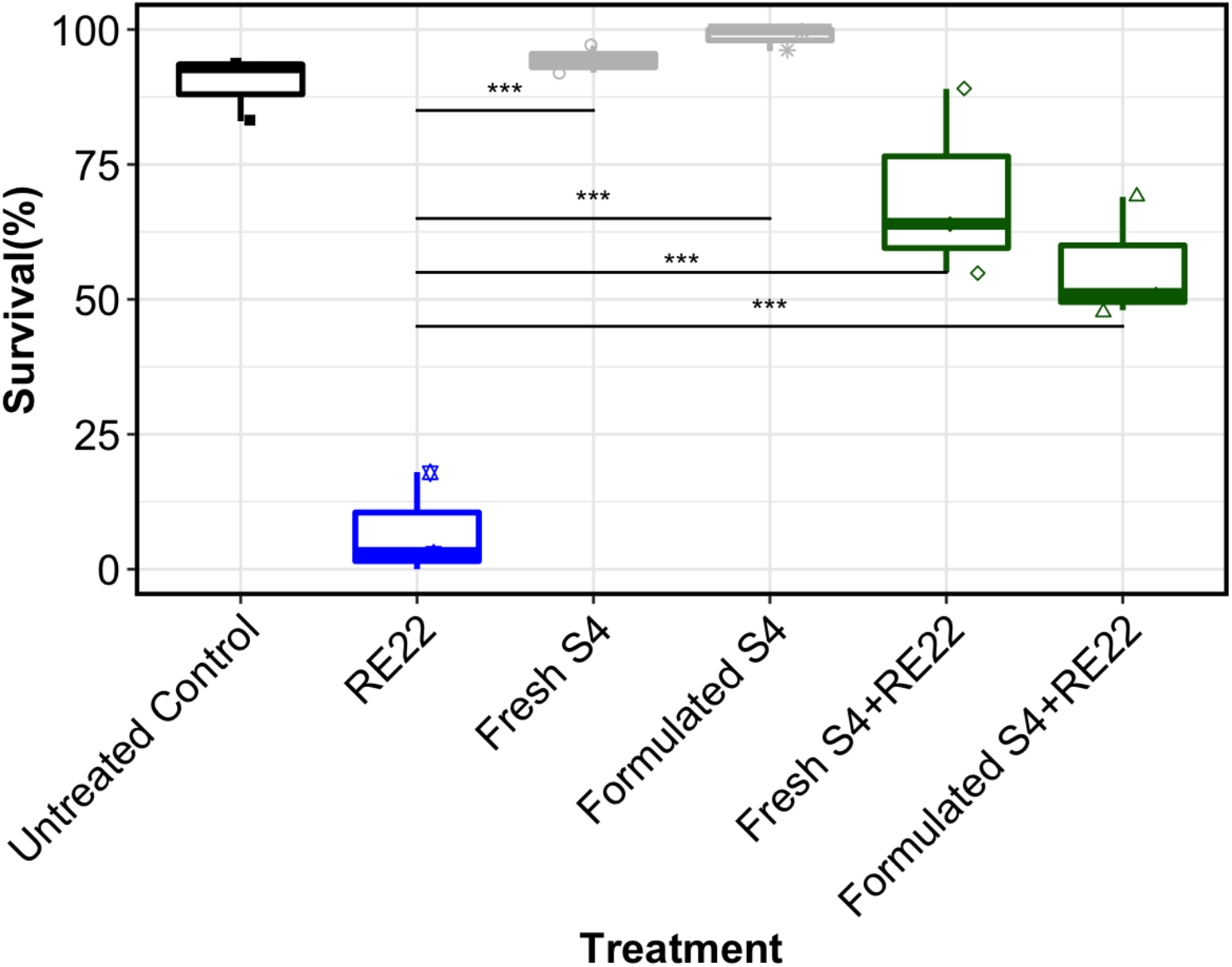
Treatment of larvae with S4 formulation in the laboratory led to increased larval survival to challenge with the pathogen RE22. Effect of pre-incubation of oyster larvae with *Phaeobacter inhibens* S4 fresh culture or formulation on survival after challenge with *Vibrio coralliilyticus* RE22 (n = 50-100 larvae per well, 3 wells per treatment). Survival was measured 24 hours after RE22 challenge (10^5^ CFU/mL) and 48 hours after addition of the probiotic S4 (10^4^ CFU/mL). Data are shown as box plots (median is shown by the line that divides the box into two parts; upper quartile - upper edge of the box - represents 75% value between the median and highest survival; lower quartile represents 25% value between the lowest and the median survival). Fresh S4: freshly cultured S4; Formulated S4: commercial batch of S4 stored in diluted mYP30 (1:1 with FSSW) at 4°C. *** indicates statistically significant differences between the treatments connected by the line (One-Way ANOVA, *p*<0.05).

### 3.3. Formulated probiotic treatment in the hatchery did not have a significant impact on larval growth and survival

Based on the protection conferred by the formulation to the bacterial challenge in the laboratory trials, the formulation (YP_SW_4) was tested in hatchery conditions. Variability in larval growth and survival between trials within a hatchery and between hatcheries was observed, with one trial (Trial 2; VIMS, July 2019) showing lower survival (*i.e*., larval crash) for both control and probiotic-treated tanks (Figure 4B). Daily treatment of larvae with the formulation in the hatcheries did not have a significant impact on larval survival or growth (specific growth rate, SGR) in any of the trials in the hatcheries, in the absence of pathogen (One-way ANOVA; *p* > 0.05; Figure 3; Figure 4).

**Figure 3.**
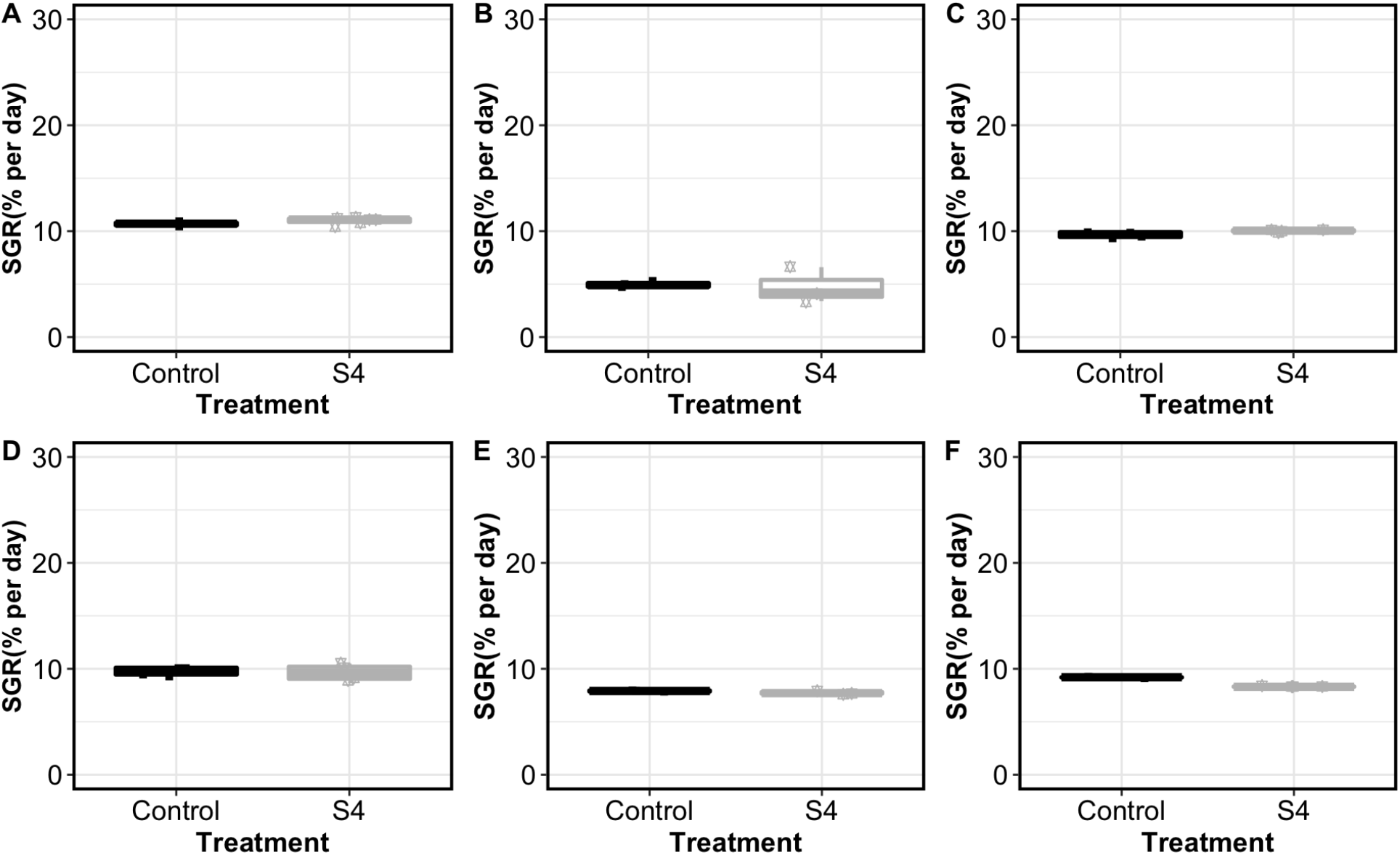
Daily treatment of eastern oyster larvae in the hatchery with S4 formulation daily did not affect larval growth. Larval oysters were treated daily with S4 at a dose of 10^4^ CFU/mL of water in the tank since day 1 post fertilization. Box plot (median, upper and lower quartile, see legend for Fig. 2) represents the data for the specific growth rate (SGR) of larval oysters at the end of each trial period (6-14 days post fertilization, n = 3 - 6 tanks per treatment, Table 1). (A) Trial 1, VIMS; (B) Trial 2, VIMS; (C) Trial 3, VIMS; (D) Trial 4, VIMS; (E) Trial 5, MOOK; (F) Trial 6, MOOK. Control: no probiotic provided; S4: *P. inhibens* S4 formulation.

**Figure 4.**
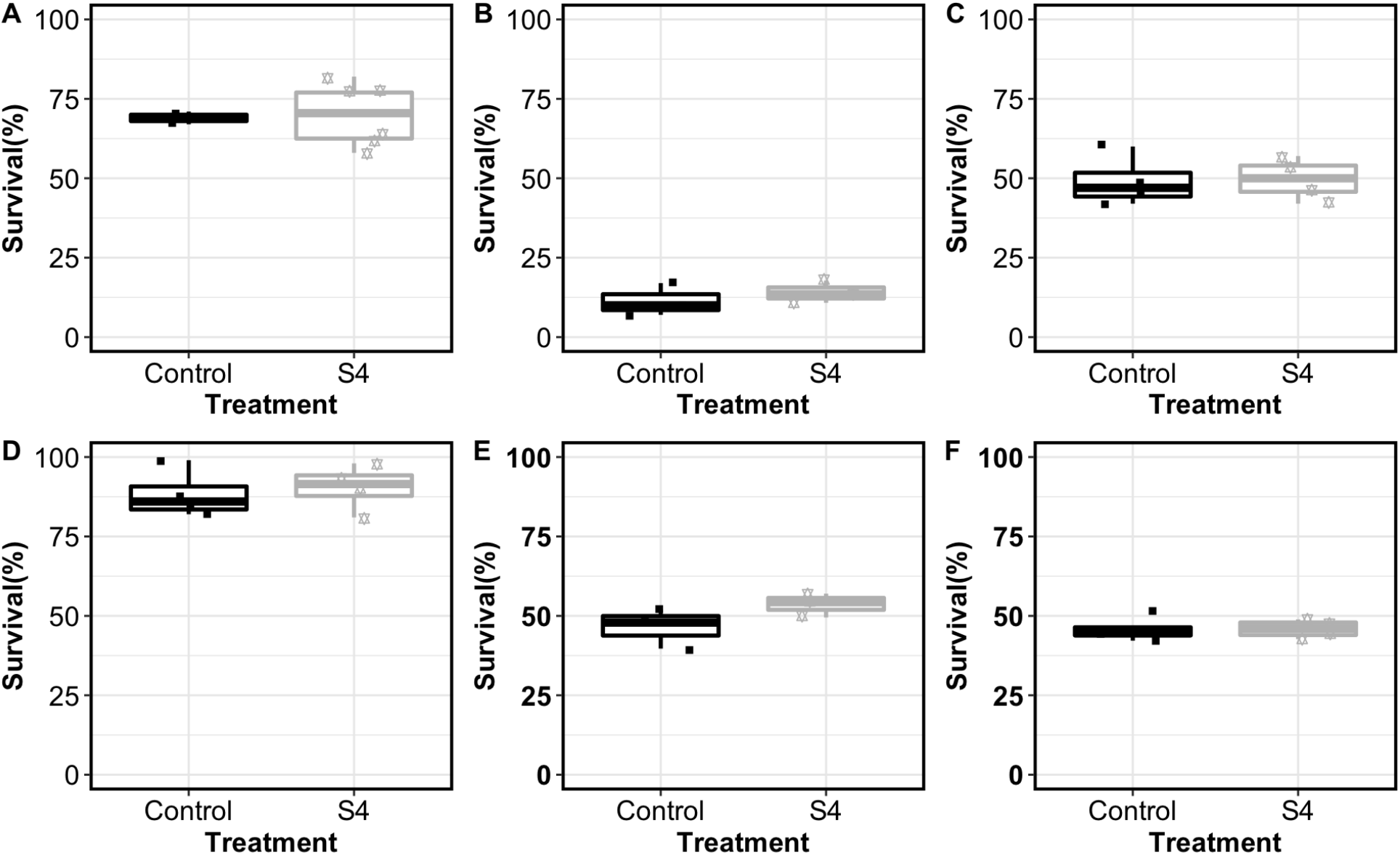
Daily treatment of eastern oyster larvae in the hatchery with S4 formulation daily did not affect survival. Larval oysters were treated daily with S4 at a dose of 10^4^ CFU/mL of water in the tank since day 1 post fertilization. Box plot (median, upper and lower quartile, see legend for Fig. 2) represents the data for larval oysters survival (in percent of total larvae stocked in tanks) at the end of each trial period (6-14 days post fertilization, n = 3 - 6 tanks per treatment, Table 1). (A) Trial 1, VIMS; (B) Trial 2, VIMS; (C) Trial 3, VIMS; (D) Trial 4, VIMS; (E) Trial 5, MOOK; (F) Trial 6, MOOK. Control: no probiotic provided; S4: *P. inhibens* S4 formulation.

### 3.4. Effect of probiotic treatment on the amount of total culturable *Vibrio* spp. in the larvae

Daily treatment of larval tanks with the probiotic formulation did not significantly decrease the total number of culturable vibrios (as detected by culture in TCBS media) in the oyster larvae compared to control treated tanks in any of the hatchery trials (*p* > 0.05 One way ANOVA for each hatchery trial; Figure 5). Variability in *Vibrio* counts between tanks within treatments and trials was observed (from 10^3^ - 10^6^ CFU/mL), likely due to handling issues and seasonal and regional differences. Culturable vibrios in water samples were below the level of detection in all trials and both hatcheries (less than 10^2^ CFU/mL, data not shown).

**Figure 5.**
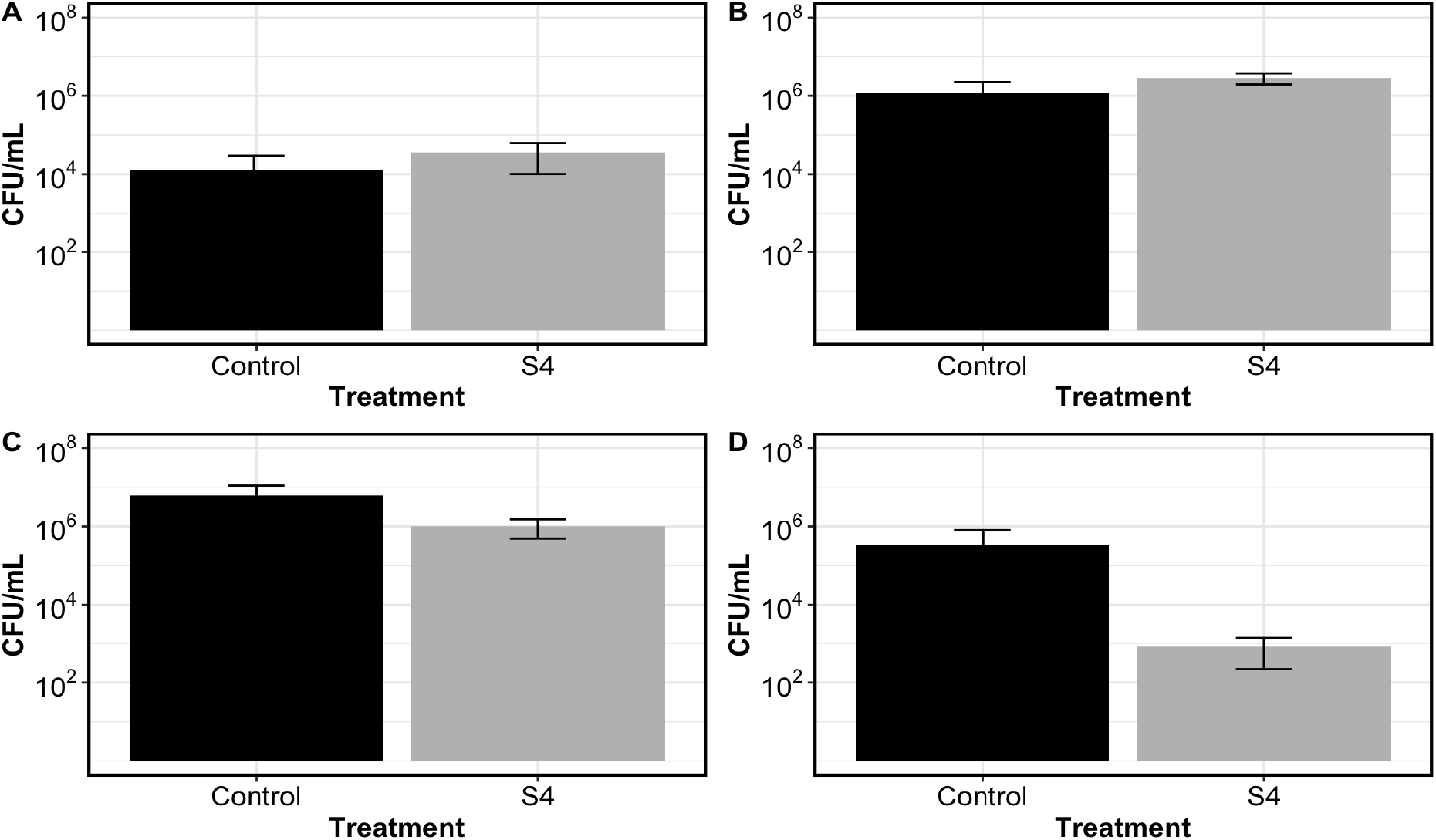
Daily treatment of eastern oyster larvae in the hatchery with S4 formulation daily did not affect total culturable vibrio levels in larvae. *Vibrio* levels (CFU/mL in samples containing around 1,000 homogenized larvae ± SD) in oyster larval samples collected from larval tanks at the hatchery were measured at the end of each trial by spot plating dilutions of homogenized larvae on TCBS agar plates. (A) Trial 2, VIMS; (B) Trial 3, VIMS; (C) Trial 4, VIMS, and (E) Trial 6, MOOK. Abbreviations: Control = no probiotic provided; S4 =*P. inhibens* S4 formulation (*p>*0.5, One Way ANOVA for each trial on log-transformed data).

### 3.5. Treatment of larvae with probiont S4 in the hatchery protected larvae from an experimental challenge with pathogen *V. coralliilyticus* RE22

Exposure of larval oysters to the S4 probiotic formulation in the hatchery significantly increased larval survival during subsequent challenge with the bacterial pathogen *V. coralliilyticus* RE22 in the laboratory (Trials 3, 4, and 6; One-way ANOVA; *p* < 0.05, Figure 6). Bacterial challenge assays for Trials 1, 2, and 5 were not performed because they were designed to confirm the safety of the probiotic formulation in each of the hatcheries before conducting further studies. In the laboratory assays, survival ranged between 72 % - 92 % for unchallenged larvae collected from both control and probiotic treated tanks. For larvae challenged with the pathogen, survival in the larvae collected from the control tanks ranged from 37 % - 41 % while survival of larvae from the probiotic-treated tanks ranged between 60 % - 76 % (relative percent survival increase of 46 % to 74 %; Table S1). Larvae treated with the S4 formulation in the hatchery demonstrated a significant increase in survival following RE22 challenge, compared to untreated oyster larvae, for all trials (One-way ANOVA for each trial, *p* < 0.05, Figure 6).

**Figure 6.**
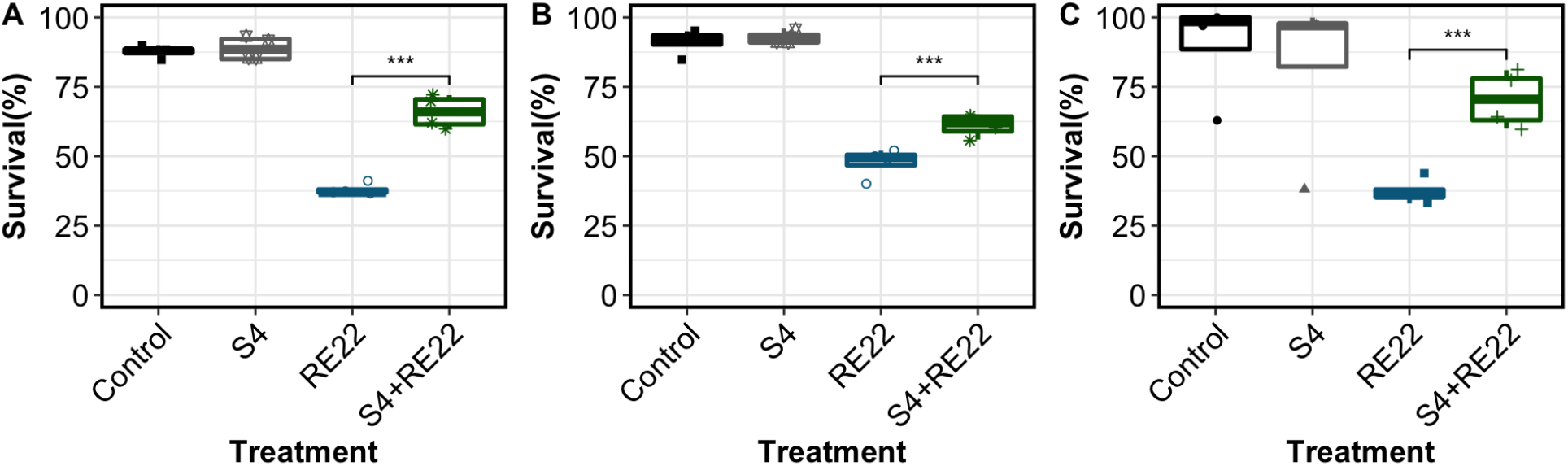
Daily treatment of eastern oyster larvae in the hatchery with S4 formulation significantly increased larval survival to pathogen RE22 challenge. Larvae from each of the hatchery tanks were transported to the laboratory at the end of each trial and exposed to a 24 hour challenge with RE22 before determining survival. Box plot (median, upper and lower quartile, see legend for Fig. 2) represents the data for percent larval survival after a 24 hour challenge with RE22 (RE22: larvae from control tanks challenged with RE22; S4+RE22: S4-treated larvae challenged with RE22) or no pathogen challenge (Control: larvae from control tanks, not challenged; S4: larvae from S4-treated tanks, not challenged). (A) Trial 3, VIMS; (B) Trial 4, VIMS, and (C) Trial 6, MOOK. S4 =*P. inhibens* S4 formulation; RE22 =*V. coralliilyticus* RE22. *** indicates statistical significance between the treatments connected by the bracket (n = 50-100 larvae per well, three wells per treatment; One-Way ANOVA for each trial, *p* < 0.05).

## 4.0. Discussion

A novel liquid formulation was developed for the commercial delivery of the gram-negative probiont S4 in oyster hatcheries. The product was found to be stable and maintained viability at 10^8^ CFU/mL or higher for a period of six weeks when stored at 4°C in an airtight container in the dark. The performance of the commercially-prepared S4 formulation tested here compared favorably to the freshly cultured S4 (as reported in [31,33]) in all laboratory experiments and hatchery trials, having no negative effect on larval growth or survival in the hatchery. As also shown previously for freshly cultured S4 [28,31], larvae treated with the probiotic formulation in the hatchery showed improved survival when experimentally challenged with the pathogen *V. coralliilyticus* RE22. This study demonstrates that the formulation is safe and effective for use in eastern oyster hatcheries to prevent larval vibriosis.

The applicability of several *P. inhibens* strains as probiotics for marine aquaculture has been assessed in several studies [25,26,27,28,29]. However, until now, no suitable formulations have been described for use in hatcheries. As a gram-negative, non-spore forming bacterium, S4 did not survive spray drying procedures commonly used to formulate gram-positive bacteria such as *Bacillus* spp. Our novel approach to formulation of this bacterium takes advantage of prior knowledge in the mechanisms allowing planktonic marine bacteria to survive in the oligotrophic conditions sometimes observed in coastal and oceanic waters [43,44]. The novelty of the formulated S4 for applications in commercial aquaculture is that it is easily delivered as a live, actively metabolizing bacteria in a liquid medium. Bacteria in the formulation remain highly viable over a period of at least 6 weeks when stored at 4°C. This formulation method differs from other commonly used techniques, such as freeze or spray drying, that put bacteria in a state of dormancy.

An effective probiotic formulation should not be harmful to the cultured larvae in the hatchery or negatively impact production. The probiotic formulation did not cause any detrimental effects to the larvae in six trials performed at two different hatcheries, confirming its safety to the larvae at the provided dose. The present study showed that there was no difference between the effect of daily treatment with fresh S4 (as reported in [31,33]) and the formulated S4 on eastern oyster larvae; *i.e*., both were safe and had no negative impact on larval growth or survival in the hatchery. Importantly, this study showed that exposure to the probiotic formulation in the hatcheries significantly improved survival of larval oysters when challenged with the pathogen RE22. This confirms results from the previous laboratory and hatchery *in vivo* challenge assays utilizing freshly cultured S4 administered prophylactically to the oyster larvae [28, 31]. Other *Phaeobacter* spp. have been shown to display a wide range of inhibitory activity against aquaculture pathogens, especially against members of the genus *Vibrio*, which are responsible for larval mortalities in aquaculture [10,45,47,48,49]. Despite showing a protective effect in oyster larvae against experimental challenge with the pathogen *V. coralliilyticus* RE22, daily treatment of larvae in the hatchery with the S4 formulation did not significantly decrease the level of culturable vibrios in the larvae in any of the hatchery trials. These results are consistent with previous studies with other *P. inhibens* strains, showing that probiotic treatment does not impact abundance of culturable vibrios in larvae [27,31,45], and suggesting that the effects of *Phaeobacter* spp. may be species-specific. Previously, microbiome analysis performed in a different study showed that probiont *Bacillus pumilus* RI0695 treatment in the hatchery leads to an increase in *Vibrio* diversity (without affecting total levels of culturable vibrios) and a shift in the composition of the *Vibrio* community to non-pathogenic species, indicating a subtle beneficial effect on larval microbial communities [46]. Probiont S4 may have direct and/or indirect effects on bacterial community diversity and composition in the hatchery due to its previously reported antibiotic, quorum quenching, and immunomodulatory actions [22,30,32]. Additional research is warranted to determine the effect of S4 on the larval microbiome, including effects on *Vibrio* spp.

Daily probiotic treatment did not significantly increase the growth or survival of larvae in any of the hatchery trials, which spanned different environmental conditions and hatchery protocols. Since larval survival was high in most of the trials, and levels of culturable *Vibrio* cells in larvae were low (*i.e*., there were no vibriosis outbreaks detected at the hatcheries), it was not expected that we would be able to detect a major effect on survival. However, S4 treatment was not able to prevent the larval crash observed in Trial 2, performed in July 2019 at VIMS, despite the consistent effect of S4 treatment protecting larvae against challenge with the bacterial pathogen RE22, and the fact that *P. inhibens* strains consistently show the ability to inhibit bacterial pathogens [10,45,47,48,49]. The lack of protection by S4 to larval losses in this trial may be due to the inability of S4 treatment to protect larvae against crashes due to causes other than vibriosis. Some eastern oyster hatcheries try to avoid spawning in July and August, since larval performance is known to be low at this time of the year due to decreased water quality (personal communications from hatchery managers) and other potential causes such as adverse environmental (*e.g*., toxins from harmful algal blooms, acifidication) or physiological (*e.g*., poor conditioning of broodstock) conditions, or other pathogens [50,51]. Also, we observed that S4 treatment did not increase the growth of the larvae in any of the trials, suggesting that probiont S4 may not provide growth enhancement benefits. Other probionts have been shown to provide direct nutritional benefits to the host through increased digestion through the release of digestive enzymes or as a direct nutritional source [22,36,52,53,54].

As seen in previous hatchery experiments with the fresh S4 culture [31], levels of variability in all the parameters that were measured (growth, survival, and culturable vibrios) were seen between tanks within treatment, between trials within a hatchery, and between hatcheries. Variability in larval performance between tanks within treatments in a trial could be due to handling and husbandry activities in the hatcheries. The frequent handling of each tank during the drain down needed to sort the larvae and maintain water quality likely led to the introduction of slightly different bacterial communities in each tank [46,55,56]. Variability in the growth and survival of the larvae between hatcheries could be due to differences in location, culture systems, water filtration methods, feeding methods, spawning events, genetic variations in broodstock and environmental conditions, to mention a few. Despite the variability in environmental conditions and performance between trials and hatcheries, there was consistency in the safety and ability to protect the larvae against RE22 pathogenic bacteria challenge when the probiotic S4 formulation was applied in the hatcheries, suggesting that S4 provides a benefit by protecting larvae against the effect of *V. coralliilyticus* RE22 infection.

## 5.0. Conclusion

This research provides evidence on the effectiveness of a newly developed approach to formulation of marine gram-negative bacteria for use as probiotics in aquaculture. This formulation approach may be useful for developing formulations of other probionts, especially gram-negative bacteria for use in marine aquaculture. The S4 formulation was shown to be safe, easy to handle, and stable to use in the hatchery environment, and it may help manage the impact of vibriosis when used prophylactically in oyster hatcheries, although it may not offer protection against other largely uncharacterized causes of larval mortality. Future research should focus on identifying the effect of S4 formulation on the microbial community of larvae, water and rearing tanks in the hatchery and combining the use of S4 with other candidate probionts or management methods to provide additional benefits to the larvae and/or prevent other causes of larval mortality.

## Acknowledgements

This work was funded by Department of Commerce/NOAA Saltonstall-Kennedy Award #NA18NMF4270193 to MGC, DNR, and DCR. ET also received support from the Blount Family Shellfish Restoration Foundation and the URI College of the Environment and Life Sciences. We are grateful to the personnel at the Aquaculture Genetics and Breeding Technology Center at Virginia Institute of Marine Science and Mook Sea Farms hatcheries, the lab of José Antonio Fernández Robledo at Bigelow Laboratory for Ocean Sciences, and undergraduate students at the University of Rhode Island Bahaa Noori and Keegan Hart for their assistance during this study. We also thank all members of the Probiotics Working Group at the University of Rhode Island.

## Data Availability Statement

The data that support the findings of this study are available from the corresponding author upon request.

## Supplementary data

**Table S1.** Effect of S4 formulation treatment in the hatchery on the ability of larvae to survive a laboratory bacterial challenge with the bacterial pathogen RE22.

| **Trial** | **Hatchery** | **Treatment** | **Relative Percent Survival (RPS, % ± SD)** |
| --- | --- | --- | --- |
| 3 | VIMS | S4 | 54 ± 11 |
| 4 | VIMS | S4 | 74 ± 11 |
| 6 | MOOK | S4 | 46 ± 19 |

Note: Relative Percent Survival (RPS) plus standard deviation (SD) of RE22-challenged oyster larvae from tanks exposed to probiotics in the hatchery relative to challenged oysters from tanks not exposed to probiotics in the hatchery. S4: S4 formulation, VIMS: Virginia Institute of Marine Science; MOOK: Mook Sea Farms.

